# Taxonomic Classification of Ants (Formicidae) from Images using Deep Learning

**DOI:** 10.1101/407452

**Authors:** Marijn J. A. Boer, Rutger A. Vos

## Abstract

The well-documented, species-rich, and diverse group of ants (Formicidae) are important ecological bioindicators for species richness, ecosystem health, and biodiversity, but ant species identification is complex and requires specific knowledge. In the past few years, insect identification from images has seen increasing interest and success, with processing speed improving and costs lowering. Here we propose deep learning (in the form of a convolutional neural network (CNN)) to classify ants at species level using AntWeb images. We used an Inception-ResNet-V2-based CNN to classify ant images, and three shot types with 10,204 images for 97 species, in addition to a multi-view approach, for training and testing the CNN while also testing a worker-only set and an AntWeb protocol-deviant test set. Top 1 accuracy reached 62% - 81%, top 3 accuracy 80% - 92%, and genus accuracy 79% - 95% on species classification for different shot type approaches. The head shot type outperformed other shot type approaches. Genus accuracy was broadly similar to top 3 accuracy. Removing reproductives from the test data improved accuracy only slightly. Accuracy on AntWeb protocol-deviant data was very low. In addition, we make recommendations for future work concerning image threshold, distribution, and quality, multi-view approaches, metadata, and on protocols; potentially leading to higher accuracy with less computational effort.

---

The family of ants (Formicidae) is a large and diverse group within the insect order, occasionally exceeding other insect groups in local diversity by far. Representing the bulk of global biodiversity (Mora et al. 2011), ants are globally found (except on Antarctica) and play important roles in a lot of ecosystems (Hölldobler et al. 1990). As ants are found to be good bioindicators, ecological and biodiversity data on them may be used to assess the state of ecosystems (Andersen 1997; Andersen et al. 2002), which is important for species conservation. Furthermore, insects are good surrogates for predicting species richness patterns in vertebrates because of their significant biomass (Andersen 1997; Moritz et al. 2001), even while using the morphospecies concept (Oliver et al. 1996; Pik et al. 1999). To understand the ecological role and biological diversity of ants, it is important to comprehend their morphology, and delimit and discriminate among species. Even working with morphospecies, a species concept is still required for identification to reach a level of precision sufficient to answer a research question. This is what is called Taxonomic Sufficiency (Ellis 1985), which must be at a certain balance or level for a research goal (Groc et al. 2010). Therefore, it is important to get a good understanding of ant taxonomy, but many difficulties arise with the complicated identification of ants to species level or to taxonomic sufficiency.

## Ant taxonomy

Classifying and identifying ant species is complex work and requires specific knowledge. While there is extensive work on this (e.g. Bolton (1994), Fisher et al. (2007), and Fisher et al. (2016)), it is still in many instances reserved to specialists. To identify ant species, taxonomists use distinct characters (e.g. antennae, hairs, carinae, thorax shape, body shininess) that differ between subfamilies, genera, and species. However, the detailed knowledge on morphological characters can sometimes make species identification difficult. Some ant species appear to be sibling species or very cryptic, and different castes complicate things further. However, with a long history on myrmecological research, ants are one of the best documented groups of insects and in recent years ant systematics have seen substantial progress (Ward 2007).

## Computer vision

In an effort to improve taxonomic identification, insect identification from images has been a subject of computer vision research in the past few years. As some early papers have shown (D. E. Guyer et al. 1986; Edwards et al. 1995; PJD Weeks et al. 1997; PJ Weeks et al. 1999; Gaston et al. 2004), a promising start has been made on automated insect identification, but there is still a long road to reaching human accuracy. Systems like a Bayes classifier (D. E. Guyer et al. 1986) or DAISY ((PJ Weeks et al. 1999) mostly utilized structures, morphometrics, and outlines. Together with conventional classifying methods (such as a principal component analysis (PCA) (P Weeks et al. 1997)) images data could be classified. Other, slightly more complex systems use simple forms of machine learning (ML) (Kang et al. 2012), such as a support vector machine (SVM) ((Yang et al. 2015) or *K*-nearest neighbors (Watson et al. 2004). An identification system for insects at the order level (including ants within the order of Hymenoptera) designed by Wang et al. (2012b), used seven geometrical features (e.g. body width) and reached 97% accuracy. Unfortunately, there are no classification studies that include ants, outside of the work of Wang et al. (2012b) on insect order level, but for other insect groups, promising results have been reported. Butterflies families (Lepidoptera) have been identified using shape, color and texture features, exploiting the so-called CBIR algorithm (Wang et al. 2012a). Insect identification to species level is harder, as some studies have shown. Javanese butterflies (Lepidoptera: Nymphalidae, Pieridae, Papilionidae, and Riodinidae) could be discriminated using the BGR-SURF algorithm with 77% accuracy (Vetter 2016). Honey bees (Hymenoptera: Apidae) could be classified with good results (*>*90%), using wing morphometrics with multivariate statistics (Francoy et al. 2008). Gerard et al. (2015) could discriminate haploid and diploid bumblebees (Hymenoptera: Apidae) based on differences in wing shape (e.g. wing venation patterns) with great success (95%). Seven owlfly species (Neuroptera: Ascalaphidae) were classified using an SVM on wing outlines (99%) (Yang et al. 2015). Five wasp species (Hymenoptera: Ichneumonidae) could be classified using PCA on wing venation data (94%) (P Weeks et al. 1997). Wen et al. (2012) classified eight insect species (Tephritidae and Tortricidae) using 54 global morphological features with 86.6% accuracy. And Kang et al. (2012) fed wing morphometrics for seven butterfly species (Lepidoptera: Nymphalidae and Papilionidae) in a simple neural network to classify, resulting in *>*86% accuracy. However, a significant disadvantage in these systems is the need for metric morphological features exploitation, which still require human expertise, supervision, and input.

## Deep learning

Deep learning (DL) may therefore be a promising taxonomic identification tool, as it does not require human supervision. DL allows a machine to learn representations of features by itself, instead of conventional methods where features need manual introduction to the machine (Bengio et al. 2012; LeCun et al. 2015). In the past few years, DL has attracted attention in research and its methods and algorithms have greatly improved, which is why its success will likely grow in the future (LeCun et al. 2015). A successful DL algorithm is the convolutional neural network (CNN), mostly used for image classification and preferably trained using GPUs. These computationally-intensive networks are designed to process (convolve) 2D data (images), using typical neural layers as convolutional and pooling layers (Krizhevsky et al. 2012; LeCun et al. 2015) and can even work with multi-view approaches (Zhao et al. 2017). A simple eight layer deep CNN has strongly outperformed conventional algorithms that needed introduced features (Held et al. 2015). It is also common practice that deep neural networks outperform shallow neural networks (Chatfield et al. 2014). In recent years, CNN technology has advanced greatly (LeCun et al. 2015; Mishkin et al. 2017; Wäldchen et al. 2018), and many biological relevant studies have shown promising results (as can be read in the next Section: Related deep learning studies).

*Related deep learning studies* CNNs have been used in plant identification (Lee et al. 2015; Lee et al. 2016; Dyrmann et al. 2016; Barré et al. 2017; Sun et al. 2017), plant disease detection (Mohanty et al. 2016) and identification of underwater fish images (Qin et al. 2016), all with high accuracy (71% – 99%). Applied examples with high accuracy include classification of different qualities of wood for industrial purposes (79%) (Affonso et al. 2017), identifying mercury stained plant specimens from non-stained (90%) (Schuettpelz et al. 2017), and identification of specimens using multiple herbariums (70% – 80%) (Carranza-Rojas et al. 2017). Especially studies like the last two are important for natural history collections, because such applications can benefit research, speed up identification and lower costs.

## Contributions

Here, we explore an alternative approach to taxonomic identification of ants based on computer vision and deep learning, using images from AntWeb (AntWeb.org 2017[a]). AntWeb is the world’s largest and leading online open database for ecological, taxonomic, and natural history information on ants. AntWeb keeps records and high quality images of specimens from all over the world, usually maintained by expert taxonomist curators. Ant mounting and photographing of specimens usually follows the AntWeb protocol (AntWeb.org 2018), which specifies standards for a dorsal, head and profile view. Considering that automating identification could greatly speed up taxonomic work and improve identification accuracy (PJD Weeks et al. 1997; Gaston et al. 2004), this work could assist in solving collection impediments. In this research, we make a start with automatic classification of ant species using images and present which shot type will classify ants the best, together with a multi-view approach. In the end we will discuss the results from different data sets and write recommendations for future work in an effort to improve taxonomic work and increase classification accuracy.

## Materials and Methods

First presented are the data sets, the process involving quality of data and, creating test sets. We used different shot types to find which type classifies best. In a different approach the three shot types are combined to one image for multi-view training. More test data for all shot types is a worker-only set and an AntWeb protocol-deviant set. Secondly, image augmentation is described and explained. Thirdly, the proposed model with its architectural decisions is discussed, and lastly the model related preprocessing actions.

### Data material

We collected all images and metadata from AntWeb (AntWeb.org 2017[a]), where images follow specific protocols for mounting and photographing with dorsal, head and profile views(AntWeb.org 2018). The intention was to work with 100 species, but the list was truncated at the 97 most imaged. This ensured the data included all species with 68 or more images, leaving out all species with 67 images or fewer. On May 15, 2018, catalog number, genus and species name, shot type, and image for imaged specimens of the 97 species were harvested from AntWeb, through its API version 2 (AntWeb.org 2017[b]). This first data set with a total of 3,437 specimens and 10,211 images is here referred to as *top97species Qmed def*. The distribution of images per species for the dorsal shot type (3,405 images), head (3,385) and profile (3,421) can be seen in Figure 1 on page 27 and Table 1 on page 34. We partitioned the images randomly in non-overlapping sets: approximately 70%, 20%, and 10% for training, validation, and testing, respectively (see Table 1 on page 34). The 70%-20%-10% was used in every consecutive dataset involving training. We downloaded images in medium quality, accountable for 233 pixels in width and ranging from 59 pixels to 428 pixels in height (for sample images see Figure 2 on page 28).

**Table 1.** Image distribution for different data sets for training, validation and test sets for 70%, 20% and 10%, respectively. top97species _Qmed_ def_ clean wtest and statia2015 rmnh have no training and validation images, because they are test data sets.

| Shot type |  | *def* | *def-clean* | *def-clean-multi* | *def-clean_wtest* | *statia2015-rmnh* |
| --- | --- | --- | --- | --- | --- | --- |
| Specimens |  | 3,437 | 3,407 | 3,322 | 2,843<sup>a</sup> | 28 |
| Total | Dorsal | 3,405 | 3,362 | - | 264 | 28 |
|  | Head | 3,385 | 3,378 | - | 279 | 28 |
|  | Profile | 3,421 | 3,377 | - | 278 | 28 |
|  | Stitched | - | - | 3,322 | - | - |
| Training | Dorsal | 2,381 | 2,354 | - | 0 | 0 |
|  | Head | 2,364 | 2,358 | - | 0 | 0 |
|  | Profile | 2,392 | 2,362 | - | 0 | 0 |
|  | Stitched | - | - | 2,322 | - | - |
| Validation | Dorsal | 691 | 681 | - | 0 | 0 |
|  | Head | 689 | 689 | - | 0 | 0 |
|  | Profile | 692 | 679 | - | 0 | 0 |
|  | Stitched | - | - | 675 | - | - |
| Test | Dorsal | 333 | 327 | - | 264 | 28 |
|  | Head | 332 | 331 | - | 279 | 28 |
|  | Profile | 337 | 336 | - | 278 | 28 |
|  | Stitched | - | - | 325 | - | - |

<sup>a</sup> 2,843 specimens were marked as valid worker specimens and, therefore, were possible specimens for the worker only test set.

**Figure 1.**
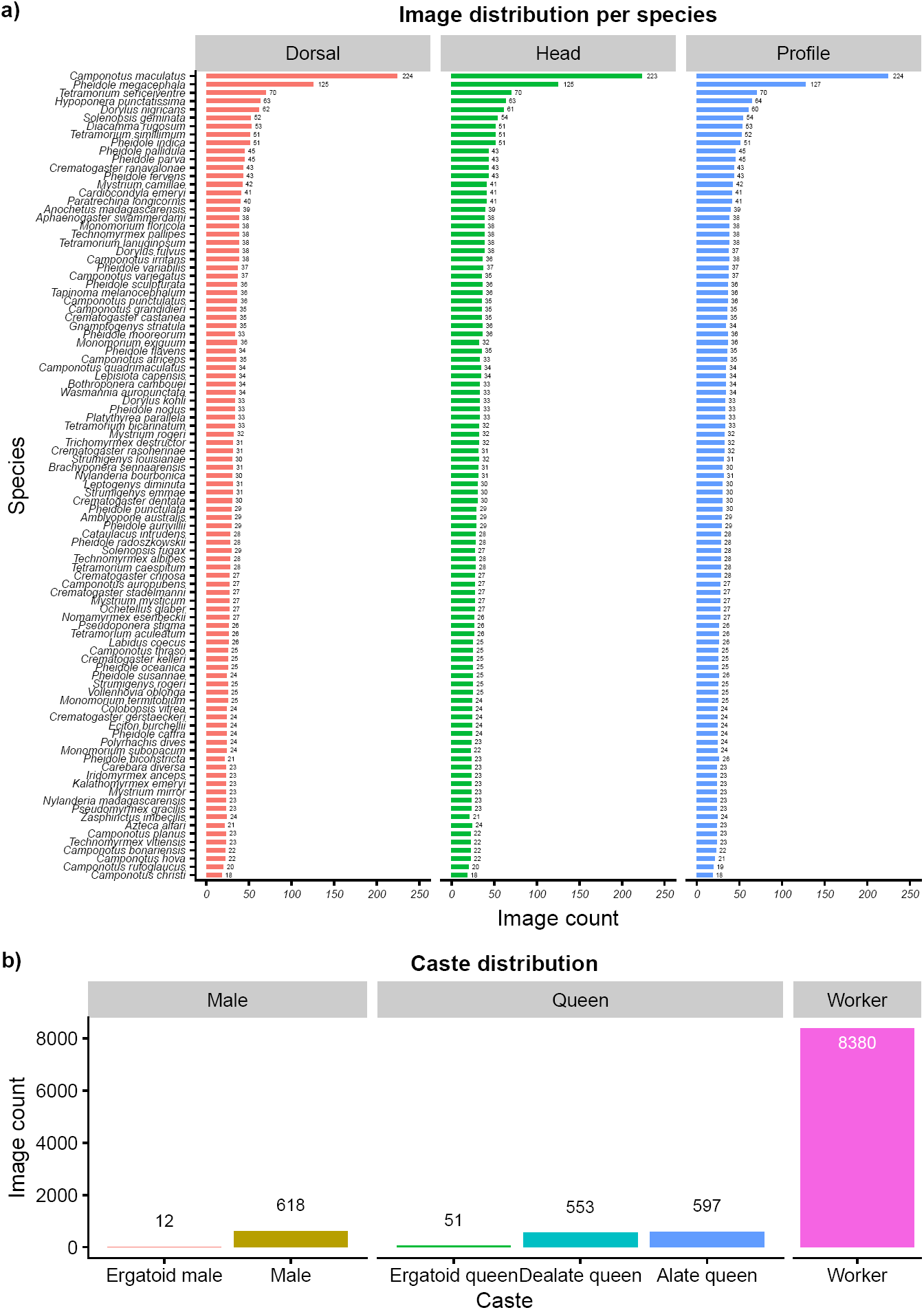
a) Histogram showing the ranked distribution for the 97 most imaged species per shot type (dorsal in red, head in green and profile in blue) for *top97species Qmed def*. Species are ranked for the combined shot type image count. Combined image counts ranges from 671 images for *Camponotus maculatus* (Fabricius, 1782) to 54 images for *Camponotus christi* (Forel, 1886). b) Histogram showing the image distribution for the different castes in *top97species Qmed def*.

**Figure 2.**
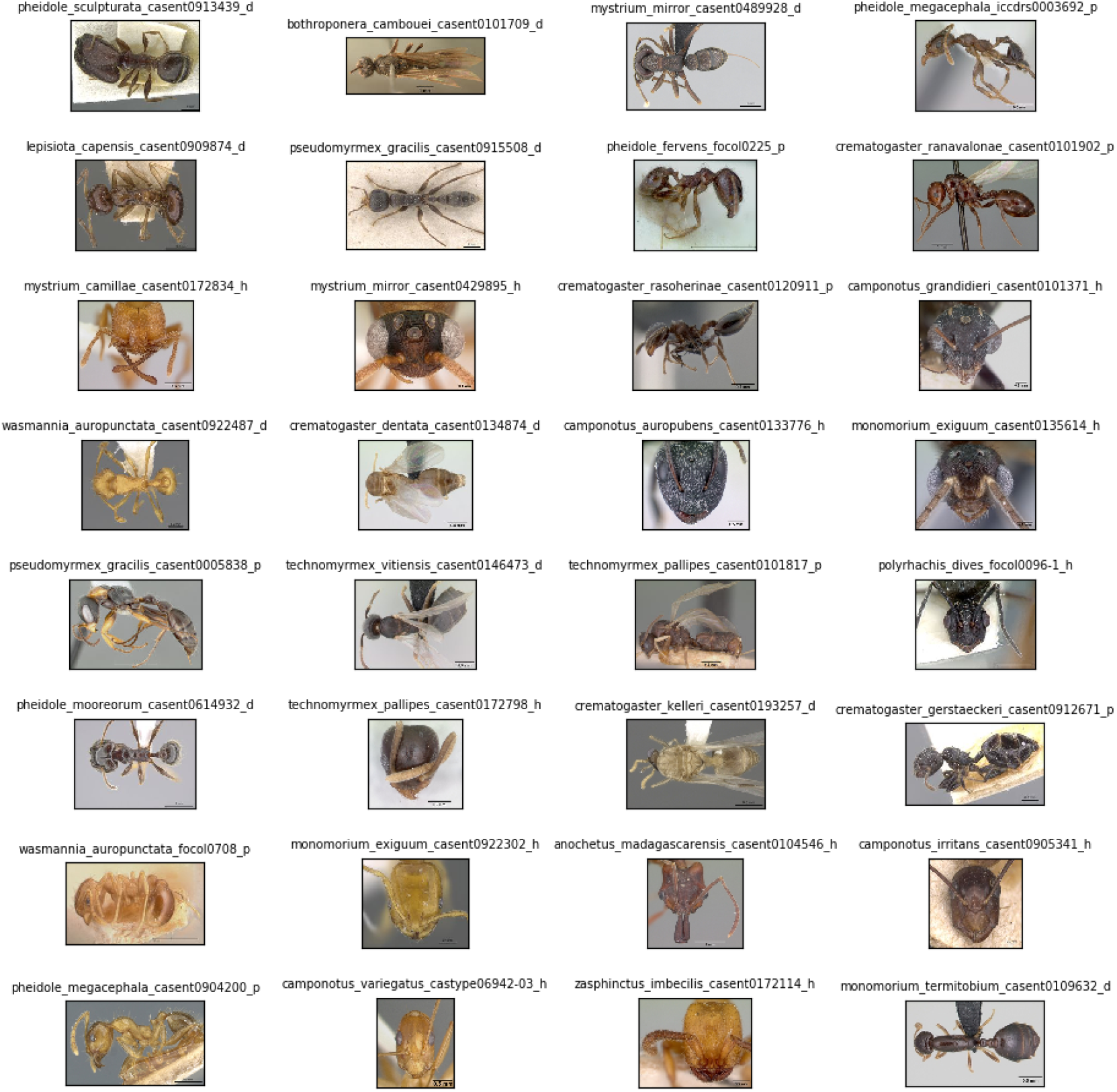
Sample images from *top97species Qmed def* showing the diversity in species, shot types, mounting, background, and specimen quality. The images have not been preprocessed. Images were downloaded from AntWeb.

#### Cleaning the data

This initial data set still contained specimens that miss a gaster and/or head or are close ups of body parts (e.g. thorax, gaster, or mandibles). A small group of other specimens showed damage by fungi or were affected by glue, dirt or other substances. These images were removed from the dataset, as these images are not representing complete ant specimens and could affect the accuracy of the model. A total of 94 images (46 specimens) were omitted from training, validation and testing (dorsal: 43, head: 7, profile: 44), resulting in 10,117 images for 3,407 specimens for a new dataset named *top97species_Qmed_def_clean*. Most of the images of detached heads could still be used, as the heads were glued on pinned paper points and looked just like non-detached head images.

#### Multi-view data set

In order to create a multi-view dataset we only included specimens in *top97species_Qmed_def_clean* with all three shot types. A total of 95 specimens (151 images) had two or fewer shot types and, thus could not be used. This list was combined with the bad specimen list for a total of 115 specimens (as there was some overlap with the one/two shot specimens and bad specimens). We removed these 115 specimens from the initial dataset so 3,322 specimens remained, all with three images per specimen per shot type, in a dataset named *top97species_Qmed_def_clean_multi* (see Table 1 on page 34). The most imaged *Camponotus maculatus* (Fabricius, 1782) had 223 three-shot specimens and the least imaged species *Camponotus christi* (Forel, 1886) only 18. Before stitching, we scaled all images to the same width, using the width of the widest image. If after scaling an image had fewer pixels in height than the largest image, black pixels were added to the bottom of this image to complement the height of the largest image (example in Figure 3 on page 29). We did not consider the black pixels as a problem for classification, because almost all stitched images had black pixel padding. The model will therefore learn that these black pixels are not representing discriminating features between species. Now, the images were combined in a horizontally stacked dorsal-head-profile image, followed by normalizing pixel values to [-1, 1] and resizing width and height to 299 *×* 299 pixels.

**Figure 3.**
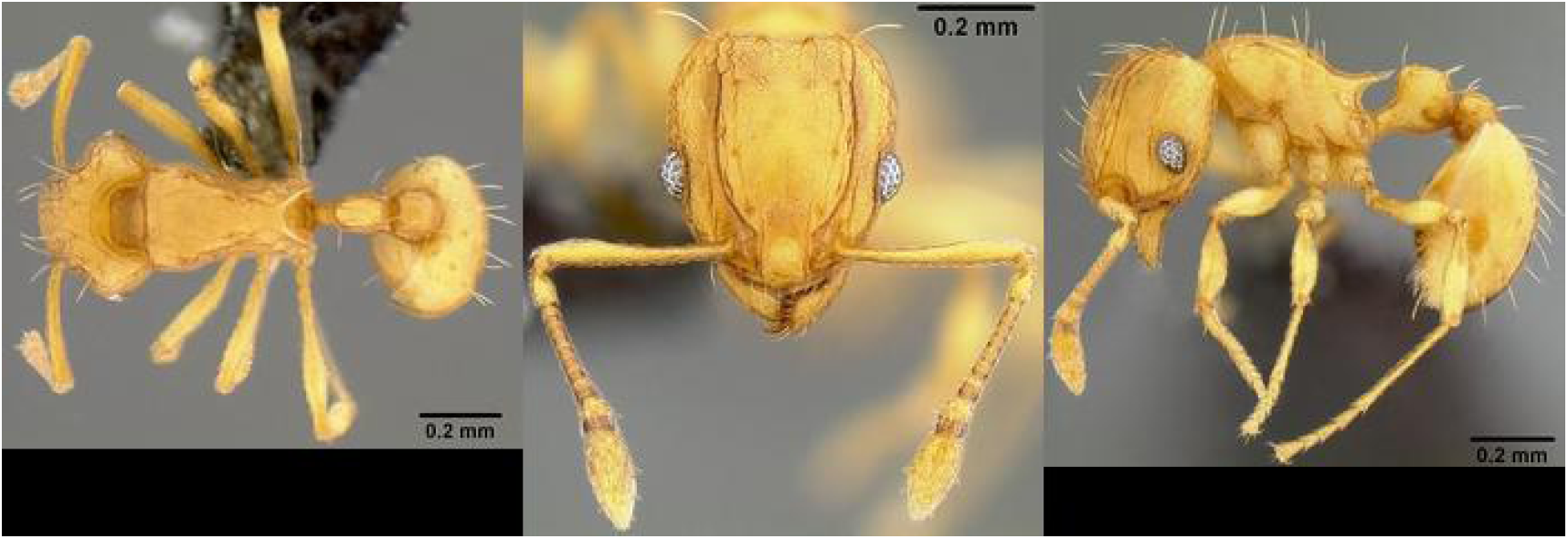
Sample image of a stitched image of the dorsal, head and, profile shot type for a *Wasmannia auropunctata* (Roger, 1863) worker (casent0171093). This image has not been preprocessed. Photo by Eli M. Sarnat / URL: https://www.AntWeb.org/specimenImages.do?name=casent0171093. Image Copyright AntWeb 2002 - 2018. Licensing: Creative Commons Attribution License.

#### Worker only test set

We labeled all specimens with their correct caste manually, as AntWebs API version 2 did not support the use of castes (support for this will be in version 3 (AntWeb.org 2017[c])). We considered alate, dealate and ergatoid queens, (ergatoid) males and intercastes as non-workers (i.e. reproductives), with no intercastes in the data set. Over 80% of *top97species_ Qmed_ def _clean* appeared to be workers (Figure1b on page 27). Consequently, 651 specimens (1,831 images) were marked as reproductives, with potential exclusion from a test set copy of *top97species _Qmed_ def _clean*. A total of 63, 52 and 58 images, for dorsal, head, profile respectively, were removed from this copy to create a test set named *top97species_ Qmed_ def _clean wtest*. The number of images in *top97species _Qmed_ def _clean wtest* set are 264, 279 and 278 for dorsal, head and profile, respectively (see Table 1 on page 34). Unfortunately, for a few species all test images were from reproductive specimens, resulting in no test images for that species. The dorsal set had five species with no test data, head only one and profile three.

#### St. Eustatius 2015 collection

In a 2015 expedition to St. Eustatius, researchers of Naturalis Biodiversity Center collected an extensive amount of flora and fauna (Andel et al. 2016). During this expedition, researchers also collected a considerable number of ant samples, now stored at Naturalis Biodiversity Center, in Leiden, the Netherlands. Most of these species all had one or more specimens imaged, and the majority of this collection was identified by expert ant taxonomists. From this collection, we extracted images of species shared with *top97species _Qmed_ def* in a new data set we refer to as *statia2015 rmnh*. This test data set of seven species with 28 images per shot type (see Table 1 on page 34) is used to assess whether the model can be applied to AntWeb protocol-deviant collections, indicating if an application will be of practical use to natural history museums and collections with existing image banks.

### Data augmentation

The issue of a small data set (*<*1 million training images) can be tackled by using image augmentation, a very common method used in DL (Krizhevsky et al. 2012). In order to artificially increase the training set, we applied label-preserving image augmentation randomly to training images during the forward pass in the training phase. Images were randomly rotated between −20° and 20°, vertically and horizontally shifted between 0% and 20% of the total height and width, horizontally sheared for maximally 20°, zoomed in for maximally 20% and horizontally flipped. It did not make sense to do heavier or other transformations, e.g. vertical flipping as ant images will never be upside down. With data augmentation, model performance is boosted because the model becomes more robust to inconsistencies in ant mounting and to within-species variation. Data augmentation can decrease the error rate between training and test accuracy, and therefore reduce overfitting (Wong et al. 2016). For data augmentation examples see Figure 4 on page 30.

**Figure 4.**
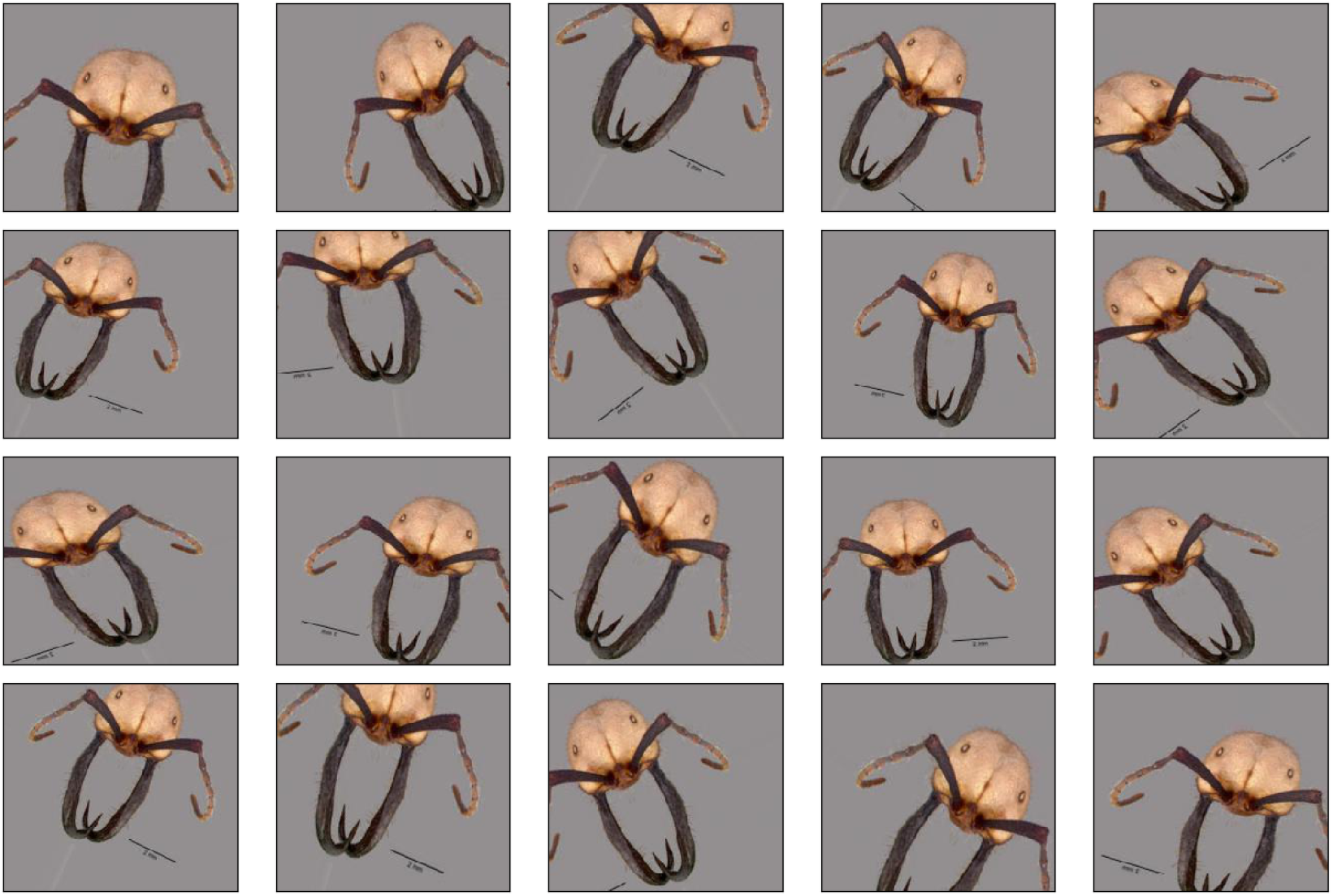
Example of random data augmentation on a medium quality head view image of a worker of *Eciton burchellii* (Westwood, 1842) (casent0009221). These images have been preprocessed and resized before augmentation. Original photo by / URL: https://www.AntWeb.org/bigPicture.do?name=casent0009221&shot=h&number=1. Image Copyright AntWeb 2002 - 2018. Licensing: Creative Commons Attribution License.

### Deep learning framework and model

We did all of the programming in Python, mostly utilizing the open source deep learning framework Keras (Chollet 2015), with the TensorFlow framework as backend (Abadi et al. 2016). We ran all experiments on a Windows 10 (64 bit) computer with a 3.50 GHz Intel Xeon E5-1650 v3 CPU and an Nvidia GeForce GTX Titan X (12GB). The network we used was Inception-ResNet-V2 (Szegedy et al. 2016) because of its efficient memory usage and computational speed. We added four top layers for this classification problem to create a modified version of Inception-ResNet-V2 (Fig 5 on page 31), in order:

**Figure 5.**
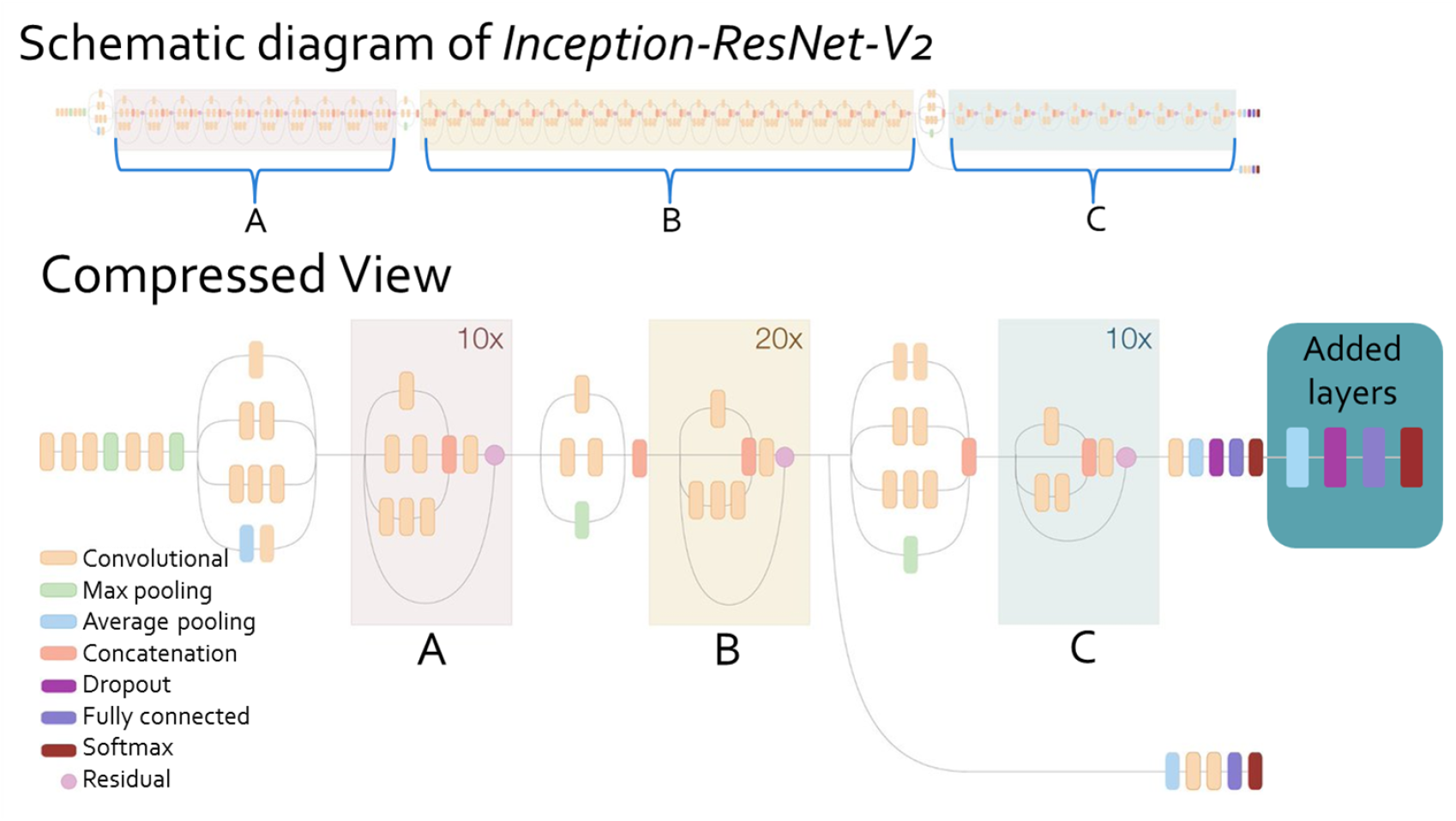
A modified version of Inception-ResNet-V2 (Szegedy et al. 2016) was used as the classifying model. It is built using 3 main building blocks (block A, B and C), each with its own repeating layers. On top of the shown network, four top layers were added, in order: global average pooling layer, dropout, fully connected layer with ReLU, and a fully connected softmax layer. Image is adjusted from: https://ai.googleblog.com/2016/08/improving-inception-and-image.html.

1. Global average pooling layer to minimize overfitting and reduce model parameters (Lin et al. 2013).

2. Dropout layer with 50% dropout probability to minimize overfitting (Srivastava et al. 2014).

3. Fully connected layer with the ReLU function as activation (Glorot et al. 2011).

4. Fully connected softmax layer to average prediction scores to a distribution over 97 classes (Krizhevsky et al. 2012).

As transfer learning is found to be a favorable method during training (Yosinski et al. 2014), we initialized with pre-trained weights (for inception models trained by Keras-team (MIT license) using the ImageNet data set (Deng et al. 2009)). We found transfer learning and fine-tuning from ImageNet to be consistently beneficial in training the ant classification models (no layers were frozen) as it greatly decreased training time. To update the parameters we used the Nadam optimizer (Dozat 2016), which is a modification of the Adam optimizer (Kingma et al. 2014) using Nesterov momentum. Nesterov momentum is usually superior to vanilla momentum (Ruder 2016), which is used in Adam. We initialized Nadam with standard Keras settings (e.g. *decay* = 0.004), except one: the learning rate was set to 0.001 and allowed to change if model improvement stagnated.

### Preprocessing

Before training, we normalized pixel values to [−1, 1] to meet the requirements of Inception-ResNet-V2 with a TensorFlow backend. Furthermore, we resized images to 299 *×* 299 pixels in width and height with the “nearest” interpolation method from the python Pillow library. We kept the images in RGB as for some specimens color could be important, giving them 3 pixels in depth. In the end, input was formed as *n* × 299 × 299 × 3 with *n* as batch number.

## Results

We configured the model to train for a maximum of 200 epochs if not stopped early. The batch size was 32 and the iterations per epoch were defined as the number of images divided by batch size, making sure the model processes all training data each epoch. We programmed the model to stop training if the model did not improve for 50 continuing epochs (due to early stopping) to prevent overfitting. Model improvement is defined as a decline in the loss function for the validation set. We programmed learning rate to decrease with a factor of approximately 0.1 if the model did not improve for 25 continuing epochs. During training, weights were saved for the best model and at the final epoch. Lastly, training, validation and test accuracy and top 3 accuracy were saved after training. Top-*n* accuracies, (commonly used with *n* = 1, 3, 5, 10), are accuracies that show if any of the *n* highest probability answers match the true label. The above settings were applied to all experiments.

### Shot type training

In all shot type experiments, validation top 1 and top 3 accuracy rapidly increased the first few epochs and after around 50 − 75 epochs the models converged to an accuracy plateau (Figure 6 on page 32). During training, the learning rate was reduced by factor 10 at epoch 47 for dorsal, epoch 66 and 99 for head, epoch 54 and 102 for profile, and epoch 50 and 80 for multi-view. At these accuracy plateaus, the models practically stopped improving, so early stopping ceased training at epoch 100, 122, 125, and 104 epochs for dorsal, head, profile, and stitched, respectively. Training usually completed in three and a half hours to four and a half hours, depending on the experiment.

**Figure 6.**
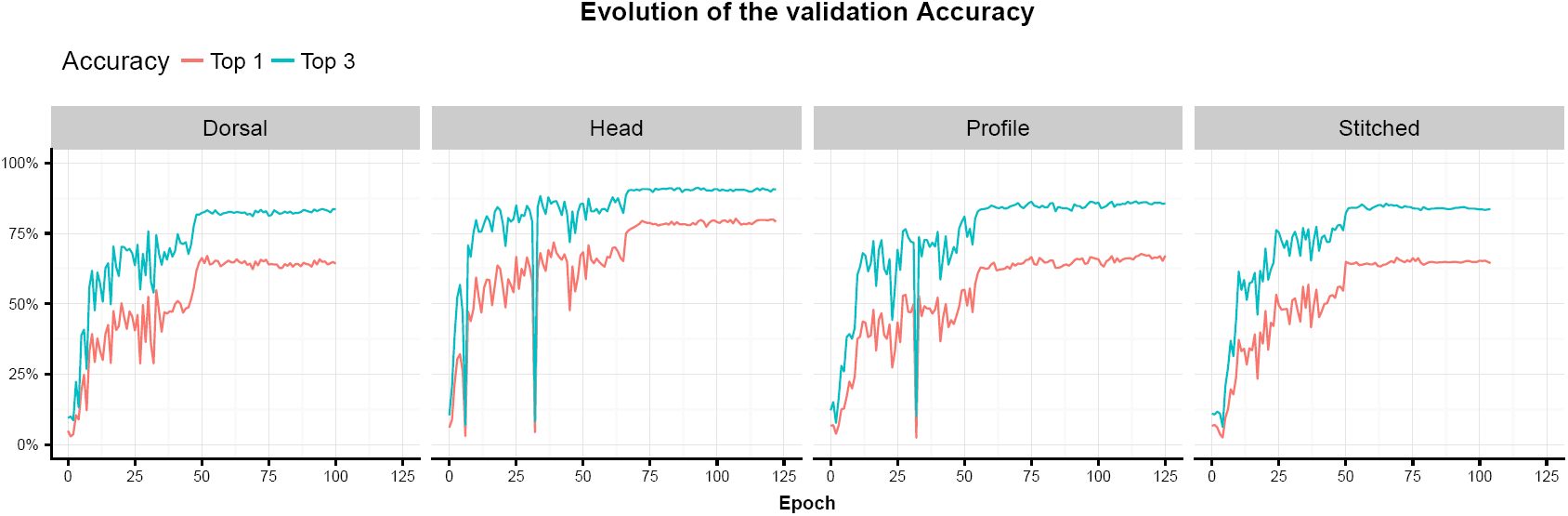
Evolution of the validation accuracy for *top97species_ Qmed_ def_ clean* for different shot types during training (in red: top 1 accuracy, in blue: top 3 accuracy). Where the line ends, training was ceased due to early stopping. From left to right: dorsal, head, profile and stitched shot type.

#### Unclean data test results

Test accuracy on *top97species_ Qmed _def* reached 65.17%, 78.82%, and 66.17% for dorsal, head, and profile views, respectively (Table 2 on page 35). Top 3 accuracy reached 82.88%, 91.27%, and 86.31% for dorsal, head, and profile view, respectively. Genus accuracy reached 82.58%, 93.98%, and 86.94% for dorsal, head, and profile view, respectively. Top 1, top 3 and genus accuracies were obtained directly after training where the model was in its validation accuracy plateau. Therefore, these accuracies do not represent the best model, of which the accuracies are shown later.

**Table 2.**
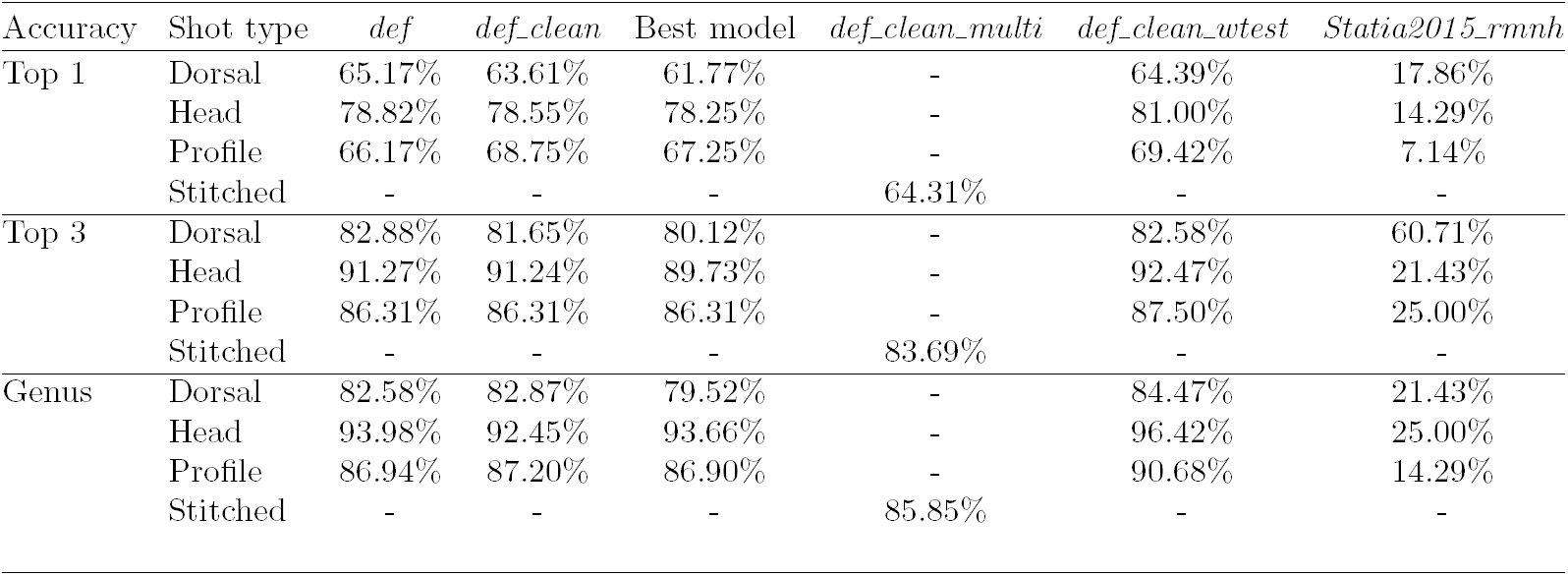
Test accuracies for different data sets and all shot types. Top 1, top 3 and genus accuracy results are shown.

#### Clean data test results

Test accuracy on *top97species_ Qmed _def_ clean* reached 63.61%, 78.55%, and 68.75% for dorsal, head and, profile views, respectively (Table 2 on page 35). Top 3 accuracy reached 81.65%, 91.24%, and 86.31% for dorsal, head, and profile view, respectively. Genus accuracy reached 82.87%, 92.45%, and 87.20% for dorsal, head, and profile view, respectively. Top 1, top 3 and genus accuracies were obtained directly after training where the model was in its validation accuracy plateau. Therefore, these accuracies do not represent the best model, of which the accuracies are shown in the section below.

During training on *top97species_ Qmed_def_ clean*, the model with the lowest validation loss function was saved at the lowest loss. This model was viewed as the best model, as the error between training and validation was at its lowest, instead of picking the model based on the validation accuracy. The lowest loss model will represent a more robust model than the previous models with higher validation loss, despite having slightly higher validation accuracy. Using the lowest loss model on the test data of *top97species_ Qmed_ def _clean*, accuracy reached 61.77%, 78.25%, and 67.26% for dorsal, head, and profile view, respectively (Table 2 on page 35). Top 3 accuracy reached 80.12%, 89.73%, and 86.31% for dorsal, head, and profile view, respectively. Genus accuracy reached 79.52%, 93.66%, and 86.90% for dorsal, head, and profile view, respectively.

Breaking down the top 1 prediction for the lowest loss models shows that most of the predictions were correct. To visualize the classification successes and errors we constructed confusion matrices using the true and predicted labels (Figure 7 on page 33). A bright yellow diagonal line indicates that most of the species were classified correctly.

**Figure 7.**
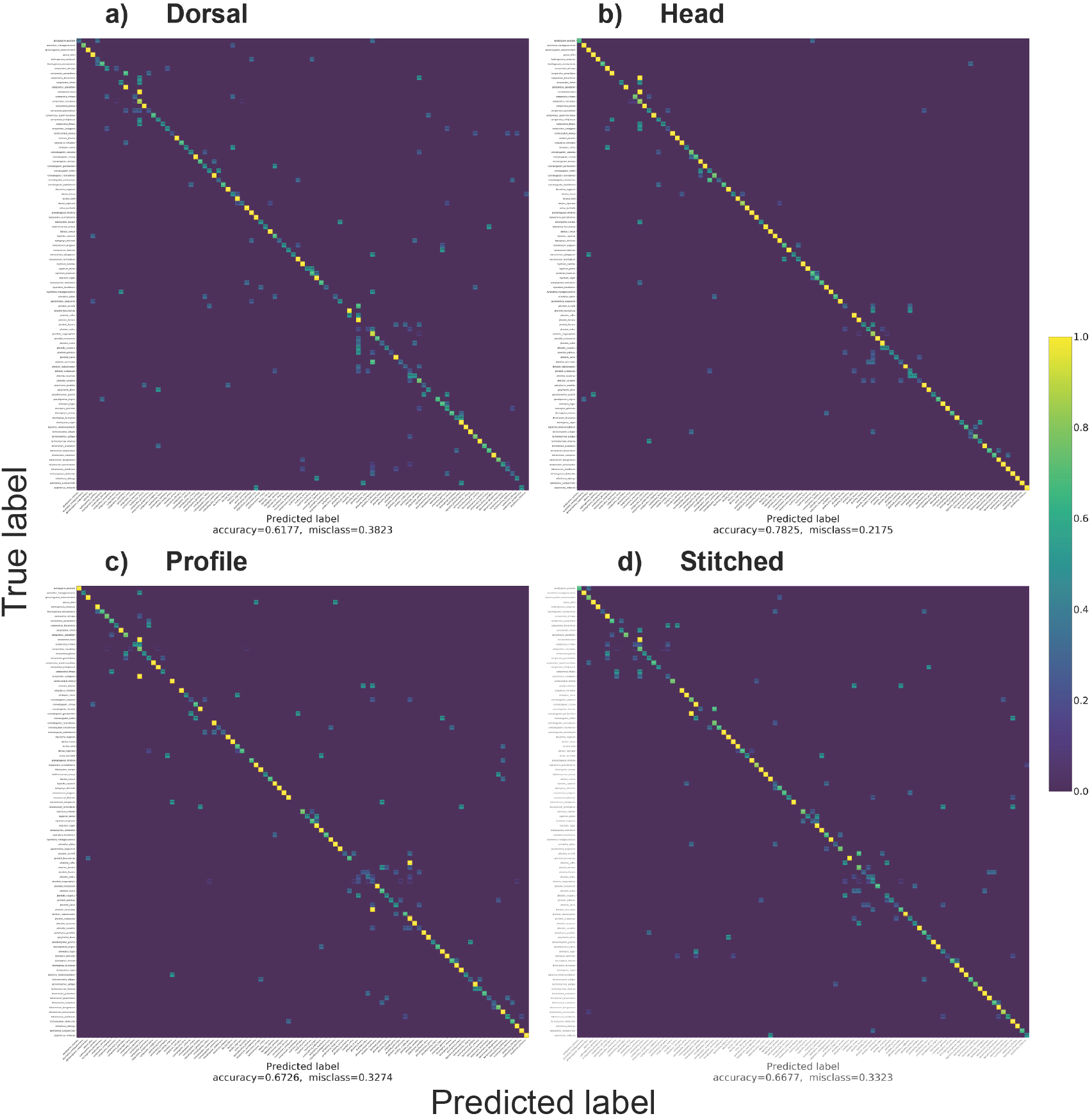
Confusion matrices showing the true label (x-axis) and predicted label (y-axis) for the dorsal (a), head (b), profile (c) and stitched (d) test sets. Each row and column represents a species. Classification accuracies are 0.6177 (a), 0.7825 (b), 0.6726 (c) and 0.6677 (d) (see also Table 2 on page 35). Most confusion was found within large genera like *Camponotus* or *Pheidole*. Confusion matrices were made using the model with the lowest validation loss trained on *top97species_ Qmed _def_ clean*. Prediction accuracy is indicated by color; from 1.0 – correct (yellow) to 0.0 – incorrect (blue). Numbers in the cells are normalized to [0, 1] to show the prediction accuracy; zeroes are not shown (best viewed on computer).

#### Multi-view test results

An accuracy of 64.31% was reached on the *top97species _Qmed _def_ clean_ multi* test set (Table 2 on page 35). Top 3 accuracy reached 83.69% and genus accuracy 85.85%. Stitched validation accuracy increased the most uniform of all shot type approaches, before reaching a plateau after roughly 50 epochs (Figure 6 on page 32).

### Worker only data results

Accuracy for *top97species _Qmed_ def_ clean _wtest* reached 64.39%, 81.00%, and 69.42% for dorsal, head, and profile views, respectively (Table 2 on page 35). Top 3 accuracy reached 82.58%, 92.47%, and 87.50% for dorsal, head, and profile view, respectively. Genus accuracy reached 84.47%, 96.42%, and 90.68% for dorsal, head, and profile view, respectively. Head genus accuracy was the highest accuracy found in all experiments.

We see that the accuracies go up, but the test set also becomes smaller. To compare this, we took worker accuracy and calculated reproductive accuracy. The head shot type reproductives reached an accuracy of 65.40%, while for workers accuracy reached 81.00%, a difference of 15.60% (Table 3 on page 36). This difference is much larger than for the other shot types; dorsal: 4.04% and profile: 4.88%.

**Table 3.** Correct and incorrect predictions, and top 1 test accuracies for workers and reproductives on top97species_ Qmed_ def_ clean. Reproductive count and accuracy is calculated from the differences in correct and incorrect predictions between top97species_ Qmed _def_ clean and top97species Qmed _def _clean _wtest. Worker accuracy is taken from Table 2 on page 35.

| Shot type | Workers |  |  | Reproductives |  |  |
| --- | --- | --- | --- | --- | --- | --- |
|  | Correct predictions | Incorrect predictions | Accuracy | Correct predictions | Incorrect predictions | Accuracy |
| Dorsal | 170 | 94 | 64.39% | 38 | 25 | 60.34% |
| Head | 226 | 53 | 81.00% | 34 | 18 | 65.40% |
| Profile | 193 | 85 | 69.42% | 38 | 20 | 65.54% |

### RMNH collection test results

Accuracy for *statia2015 rmnh* reached 17.86%, 14.29%, and 7.14% for dorsal, head, and profile views, respectively (Table 2 on page 35). Top 3 accuracy reached 60.71%, 21.43%, and 25.00% for dorsal, head, and profile view, respectively. Genus accuracy reached 21.43%, 25.00%, and 14.29% for dorsal, head, and profile view, respectively. This is the only case where genus accuracy is substantially lower than the top 3 accuracy. Profile top 1 accuracy was the lowest accuracy found in all experiments.

## Discussion

We present an image-based ant classification method with 61.77% – 81.00% accuracy for different shot types. We processed the input for training in different ways and with test data including a worker-only and an AntWeb protocol-deviant test set. Consistently throughout our experiments, shot type accuracies were found to rank from low to high accuracy in the same order: dorsal *→* profile *→* head. The head shot type predominantly outperformed dorsal, profile, and stitched in accuracy by about ten percentage points most of the time, perhaps due to the fact that this shot type is more protocol stable. An additional explanation may be that discriminating characters are more concentrated in the head in some ant groups. The combined, stitched image view did not greatly increase accuracy, as the head shot type outperformed the stitched view by 6.04% – 7.58%. A not so much curious result, as the combination of multiple views in one image is the most naive way of approaching a multi-view learning problem (Zhao et al. 2017). Other approaches on a multi-view problem (discussed in Section: Recommendations for future work) would most probably have higher accuracies. Genus accuracy reached 79.52% – 96.42%, which is approximately as accurate as the top 3 accuracy (80.12% – 92.47%), sometimes slightly above it. It is, however, important to note that the CNN has no understanding of what a genus is, because it selects the label *genus species* from among a flat list. Top 3 accuracy is preferred over genus accuracy as this will show only three options, of which one is correct, where a genus accuracy could still have over 20 potential species.

Looking at the confusion matrices (Figure 7 on page 33) outliers can best be explained as specimens that are morphological-wise very comparable. This is especially the case in *Camponotus, Crematogaster* or *Pheidole*, which have a lot of species in the dataset (14, 8, and 17, respectively). In contrast, just eight other genera have two to six genera in the dataset and the rest only one. And because the species in the confusion matrices are alphabetically sorted on genus, false predictions near the yellow diagonal line are most of the time found within the correct genus for these three big genera. Therefore we speculate that inter-genus features are better distinguished than intra-genus features.

Because the majority of specimens are workers, there is most probably a bias in learning the workers from a species. We therefore speculate that the model has acquired an improved understanding and representation of workers. However, accuracy for workers did increase only slightly, when reproductives were removed from the test set. We see a slight increase in dorsal and profile worker accuracy over reproductives accuracy, but the increase is small. The only noticeable and interesting increase is for the head shot type, where workers were classified 15.60% more accurate (Table 3 on page 36). We still see a slight increase in dorsal and profile worker accuracy over reproductives accuracy, but the increase is small. It seems that discriminating workers from reproductives is best performed using the head. This could have something to do with ocelli, only present on heads of reproductives, causing trouble.

The image number threshold for the species in this data set was 68 images, which is approximately 23 images per shot type. That accounts for 16 images in the training set, which nonetheless achieved good accuracy. This means that the threshold could potentially be lower, and thus more species (with fewer than 68 images) could be incorporated. However, more species (classes) will also complicate training and test accuracy.

One of the biggest improvements in accuracy can be made by increasing the data and thus reducing variance (training error) and overfitting. The current data shows a much skewed, long tailed, distribution with the first two species containing over 10% of the total number of images. Furthermore, only *C. maculatus* and *Pheidole megacephala* (Fabricius, 1793) had over 100 stitched images out of 3,322 in total. Also important when expanding the image set is adding male and queen specimens so the classifier has improved learning of these castes. Despite the fact that Bolton (2003) provided the first big overview for male ant taxonomy, at this moment 22% of extant species still have their male castes unknown, because males are usually only found incidentally. As males have been found to be important factors in a colony and not just sperm vessels (Shik et al. 2012), it is important to include these underrepresented specimens in automatic ant identification.

Results are not shown, but species in a species complex (i.e. species with subspecies) did not complicate training and did not cause accuracy problems. This was measured using the *F*_1_-score, calculated as the harmonic mean of precision and recall. With an increasing number of species in a complex, the *F*_1_-score did not increase or decrease significantly; variation in data could not be explained by the linear relation.

Of interesting note is the labeling of this data set, as this was not managed by the author. Identifications and labels were directly taken from AntWeb, assuming that they were correct. However, there is always a chance that identifications are less accurate and certain as expected (e.g. Boer (2016)), despite being a by-expert-labeled data set. Reality is that ant identification is more complex work than labeling a cat and dog dataset for example.

Despite some obstacles and points for improvement, we have shown that processing data in different ways influences test results in different ways. In this article we demonstrated that it is possible to classify ant species from images with decent accuracy.

### Recommendations for future work

To the best of our knowledge, this is the first time ants were classified to species level using computer vision, which also means that there is a lot to improve. In this section we will discuss some possible improvements for future research in the form of recommendations.

To start, focus should lie on creating benchmark data set that is easy to enlarge and improve. To do that, first it is important to find the image threshold for the model to learn a species, which could differ per genera and species. Finding this number would shift the focus to photographing species below the threshold in reaching the threshold. To also increase the data set in the near future, specimens from natural history museums ant collections should be photographed, as it would be less time and cost expensive than collecting new specimens. These existing specimens are most likely already following AntWeb mounting standards. Hopefully this could also solve the skewed image distribution and add more three shot type specimens. In the end, this data set could serve as benchmark data for automated insect identification, and then research focus can shift to accuracy-improving efforts.

One of these efforts could be the incorporation of a hierarchical system, where the model classifies on different taxonomic levels as Pereira et al. (2016) did with orchids (Asparagales: Orchidaceae). In an effort to do this, one could do this in a series of multiple CNNs (e.g. first subfamily, then genus, then species), but also in three parallel CNNs, learning simultaneously. However, for this we first need to work on a (phylogenetic) tree and molecular data, which is a different study itself. Moreover, there is also the option to classify on caste, before classifying species, using a caste-trained CNN, and then make use of specialized workers, males and queen trained CNNs.

An other option is to incorporate metadata; e.g. biogeographic region, caste, country, collection locality coordinates, or even body size (using the included scale bar on images). Metadata could be very important, especially for species that are endemic to a specific region. Metadata could provide underlying knowledge of the characteristics. Most of this information is already present on AntWeb and ready for use.

In order to improve the multi-view approach, multiple solutions have been tried (Zhao et al. 2017). A first option is to try is using just one CNN with all images as input and with the addition of catalog number as a label will. The next option could be to train three shot type CNNs parallel and combine the output. The output can be processed as the average of three shot type predictions, or by using the highest prediction. It is also possible to overlay the three images and take the average pixel values in order to create an average single input image of a specimen.

Furthermore, as results have shown, it is very important to have the same mounting procedure and photographing protocol to get a uniform set of images. Difference in dried and alcohol material is most likely very important, but other details like type of glue, background, and zoom could potentially be important and will have to be standardized. Also to get high-detail images, the use of good image stacking software and high-resolution cameras is very important. Therefore, the recommendation is to follow the, already widely used, AntWeb protocol (AntWeb.org 2018).

In the end, research like this could assist taxonomists, natural history museums, and researchers to achieve higher taxonomic completeness, better collections and therefore improve research. But for general use the code should further be developed in an easy to use application. A functioning application with high accuracy could reduce costs, time, and energy during everyday identification work (Gaston et al. 2004). However, bear in mind that an application like this is not aimed for use in the field and there is still skill required in collecting, mounting and photographing specimens. Nonetheless, we would like to argue that automated species identification from images using a CNN has high potential.

Research in this subject should be continued, and even though DL still has some obstacles to overcome (Marcus 2018), it has already advanced a lot (Guo et al. 2016; Wäldchen et al. 2018).

## Acknowledgements

We thank AntWeb and its community for providing the necessary data, images and information to make this research possible. We would also like to thank Laurens Hogeweg for all the help and discussions on neural networks and the programming that goes with it. Furthermore, we would like to thank dr. Martin Rücklin for use of computers and guidance in the computer lab.

## Supplementary Material

Programming code and documentation is available for open access (MIT licensed) and published on URL: github.com/naturalis/FormicID.

Data available from the figshare repository: https://doi.org/10.6084/m9.figshare.6791636.v4

